# Biodegradable Nanoparticle-in-Implant Platform for Sustained and Light-Boosted Pirfenidone Delivery

**DOI:** 10.64898/2026.06.17.732517

**Authors:** Joseph Mwaniki, James Kelley, Yoonjee Park

## Abstract

Vocal fold (VF) fibrosis is a major cause of persistent dysphonia due to excessive extracellular matrix deposition and tissue stiffening that disrupt normal vocal fold vibration. Current treatment approaches are limited by the need for repeated local injections and inadequate long-term therapeutic control. Pirfenidone (PFD), an FDA-approved antifibrotic agent, has demonstrated potential for reducing fibrosis; however, its short half-life and systemic adverse effects limit conventional administration strategies. In this study, we developed a sustained and near-infrared (NIR)-responsive local delivery platform by integrating PFD-loaded poly(lactic-co-glycolic acid) (PLGA) nanoparticles into biodegradable PLGA implants for dose-controllable antifibrotic delivery.

PFD-loaded PLGA nanoparticles were fabricated using an oil-in-water emulsion solvent evaporation method and characterized by dynamic light scattering (DLS), transmission electron microscopy (TEM), and scanning electron microscopy (SEM). Nanoparticles with small, medium, and large hydrodynamic diameters were generated to evaluate the effect of particle size on release behavior. Gold nanorods (AuNRs) were incorporated to enable photothermal NIR-triggered release enhancement. The nanoparticles were subsequently loaded into non-porous PLGA (90:10) implants and evaluated for long-term *in vitro* release under physiological conditions with and without pulsed 1064 nm laser irradiation.

The nanoparticle-loaded implants demonstrated sustained PFD release for over 190 days with minimal initial burst release (<2.5%). NIR irradiation enhanced PFD release compared with non-irradiated controls across all nanoparticle sizes. Smaller nanoparticles produced greater cumulative release than medium and large nanoparticles due to shorter diffusion pathways and larger surface-area-to-volume ratios. Prior to implant fracture, cumulative PFD release reached approximately 20.2%, 14.8%, and 12.3% of total loading for small, medium, and large nanoparticle groups under 2-min irradiation conditions, respectively. Dialysis membrane studies further demonstrated that the PLGA capsule acted as an additional diffusion barrier that substantially prolonged release compared with nanoparticles alone.

Overall, this study demonstrates a hybrid nanoparticle-in-implant strategy capable of providing sustained and irradiation-enhanced local PFD delivery with tunable release characteristics. These findings support the potential of biodegradable, dose-controllable implant systems for long-term management of vocal fold fibrosis while reducing the need for repeated interventions.

## 1. INTRODUCTION

The human voice is essential for communication, and phonation occurs when airflow drives vibration of the vocal folds (Elibol et al., 2021; Miura et al., 2024). Disruption of this vibratory function can result in dysphonia and dyspnea in severe cases (Graupp et al., 2016). Persistent voice impairment can substantially affect quality of life, particularly for individuals with voice-dependent occupations (Coburn et al., 2022; Nakamura et al., 2021). Because voice production depends on precise, repeatable vibration, small changes in vocal fold structure can lead to noticeable changes in voice quality. Vocal fold scarring/fibrosis is a challenging outcome of laryngeal injury because scar-driven remodeling can permanently alter tissue biomechanics and degrade vocal function and hinder rehabilitation (Mora-Navarro et al., 2026). This is partly because fibrosis disrupts the specialized layered microarchitecture—particularly within the lamina propria/superficial lamina propria—that is required for normal vocal fold vibration (Allen, 2021; Shin et al., 2024). VF fibrosis frequently occurs following phonosurgery, mechanical/iatrogenic trauma, endotracheal intubation-related injury, and radiation therapy (Mora-Navarro et al., 2026; Zheng et al., 2023). Consequently, once the tissue’s mechanical properties shift, restoring normal vibration becomes inherently difficult. Biologically, fibrotic remodeling is associated with excessive extracellular matrix (ECM) deposition that increases tissue stiffness, reduces pliability, and impairs mucosal wave propagation—changes that directly undermine normal VF vibration and present clinically as dysphonia/hoarseness and breathiness (Gracioso Martins et al., 2022; Mora-Navarro et al., 2022; Shin et al., 2024). This is why durable therapies must ultimately address tissue mechanics, not only short-term clinical symptom relief. Despite ongoing clinical and research efforts, effective long-term therapies that reliably restore VF pliability remain limited, and there are no FDA-approved treatments that specifically target VF fibrosis (Allen, 2021; Mora-Navarro et al., 2026).

Pirfenidone (5-methyl-1-phenyl-2-[1H]-pyridone) is an oral antifibrotic agent approved by the U.S. Food and Drug Administration (2014) for the treatment of idiopathic pulmonary fibrosis (U.S. Food and Drug Administration, 2014). Beyond pulmonary fibrosis, pirfenidone has demonstrated antifibrotic activity in diverse tissues, including cutaneous scars, vocal fold scarring, abdominal adhesions, intestinal and esophageal strictures, and benign biliary strictures (Betancourt-Vicencio et al., 2022; Hirano et al., 2022; Knighton et al., 2025a; Kodama et al., 2018a; Latella & Viscido, 2020; Orozco-Perez et al., 2015; Park et al., 2021; Q. Yang et al., 2018). Although its precise mechanism of action is not fully defined, pirfenidone appears to act through dual pathways: it downregulates pro-inflammatory cytokines such as TNF-α, interleukin-1β (IL-1β) and interleukin-6 (IL-6), dampening the early injury-driven inflammatory response, and it is reported to inhibit transforming growth factor-β (TGF-β) production and TGF-β–mediated collagen synthesis, while suppressing fibroblast proliferation and myofibroblast differentiation (Antar et al., 2022; Ballester et al., 2020; S. Liu et al., 2021; Shah et al., 2021; Torre et al., 2024a). Additionally, pirfenidone has been reported to limit collagen I fibril formation and reduce the deposition of extracellular matrix components (Aimo et al., 2022; Bahudhanapati & Kass, 2017). Collectively, these anti-inflammatory and antifibrotic effects make pirfenidone a rational candidate for preventing or attenuating VF scarring. However, oral delivery has practical limitations: the drug exhibits a short plasma half-life of ∼ 3 hours, necessitating frequent dosing three times daily (2403 mg/day total dose) to maintain therapeutic levels (Aimo et al., 2022; U.S. Food and Drug Administration, 2014). This high-frequency dosing regimen, combined with systemic side effects including gastrointestinal disturbances (nausea, dyspepsia, diarrhea), photosensitivity reactions, rash, and hepatotoxicity, contributes to poor patient compliance and frequent treatment discontinuations (Han et al., 2024; Lancaster et al., 2016). These limitations underscore the need for alternative delivery strategies that can provide sustained local drug concentrations while minimizing systemic exposure and improving patient adherence.

Because the vocal folds are a highly specialized, layered tissue whose normal function depends on finely tuned vibration, they are particularly vulnerable to injury and biomechanical disruption (Manick et al., 2025). In practice, repeated local drug delivery to the vocal folds is not straightforward because access typically requires procedure-based laryngeal visualization and/or targeted injection approaches (Bar et al., 2024). Consistent with this, common procedural approaches require skilled laryngoscopic guidance and may necessitate repeat procedures due to variable patient response and/or material resorption (Brown et al., 2024; Kim et al., 2024).

Steroid injections can improve function in the short-term but often require repeat dosing, which is constrained by VF access and carries risks such as atrophy and hematoma (Axiotakis et al., 2023; Jeong et al., 2025; Wang et al., 2017; Young et al., 2016). By contrast, systemic administration is often suboptimal for a focal laryngeal target because drug distributes broadly (increasing off-target exposure) and dosing is constrained by systemic tolerability (Budală et al., 2023). Similarly, systemic or inhaled steroid therapy is often less effective for VF scarring and is prone to systemic side effects(Amin et al., 2019; Govil et al., 2014). Localized delivery is therefore attractive because it can increase drug concentration at the target site, reduce systemic side effects, and extend therapeutic exposure with minimal repeat intervention, which are widely recognized advantages of local drug delivery over systemic dosing (Ezike et al., 2023; Saberian et al., 2024; Zhu et al., 2025). Taken together, these limitations support the need for long-lasting, safe, and controllable treatment strategies for VF scarring.

Biodegradable polymeric nanoparticles—especially PLGA—are widely used as drug carriers because they are biocompatible, biodegradable, and well-suited for controlled release (Nath et al., 2024). More broadly, biodegradable polymeric nanoparticles can encapsulate diverse therapeutics (including small molecules, proteins, and nucleic acids), supporting adaptable nanoformulations across multiple delivery needs (Stevanović et al., 2025). Among biodegradable polymers, PLGA is a leading example with EMA and FDA approval and an extensive clinical track record that supports its reliability for drug delivery applications (J. Yang et al., 2024).

Encapsulation within PLGA nanoparticles can protect drug cargo from premature degradation and support tunable, sustained release, with kinetics adjusted through formulation variables such as polymer composition (lactide:glycolide ratio), molecular weight, end-group chemistry, and particle size (Bassand et al., 2022; Nath et al., 2024; Stevanović et al., 2025; J. Yang et al., 2024). In addition, PLGA undergoes hydrolytic degradation to lactic and glycolic acids that are naturally metabolized, helping avoid toxic accumulation and supporting predictable clearance after delivery (Ulianova et al., 2023). In line with this rationale, pirfenidone (PFD) nanoformulations have demonstrated feasibility and therapeutic promise in pulmonary models, including inhalable PFD-loaded nanoparticle systems designed for sustained lung delivery (Dudhat & Patel, 2022). For example, PFD-loaded chitosan nanoparticles formulated for pulmonary delivery have been reported to provide sustained release and effective lung deposition, consistent with enhancing local exposure while minimizing systemic burden (Dudhat & Patel, 2022). Similarly, PFD-loaded nanostructured lipid carriers have been reported to improve encapsulation efficiency and enable controlled release kinetics in pulmonary fibrosis models (Chettupalli et al., 2025). In dermatological/scar models, PFD has been incorporated into nanocarrier/wound-dressing platforms, where localized delivery has been reported to support wound repair and reduce scar formation, consistent with antifibrotic activity in dermal settings (He et al., 2022; Knighton et al., 2025b). Additionally, topical PFD formulations incorporating nanocarriers have also demonstrated efficacy in reducing skin fibrosis and improving wound healing in animal models, with favorable safety profiles (Torre et al., 2024b). Collectively, these preclinical findings support the rationale for developing PFD-loaded PLGA nanoparticles as a sustained-release platform for localized antifibrotic therapy in vocal fold fibrosis.

Biocompatible implantable depots are an attractive strategy for vocal fold therapy because they can anchor the drug at the target site and support localized administration that maximizes on-site exposure while reducing systemic exposure (Magill et al., 2023). This approach is particularly relevant because local anatomy and physiology can limit drug residence and exposure, and conventional formulations often struggle to maintain sufficient drug levels at the intended site over extended periods (Bhattacharjee et al., 2019; Kumar et al., 2022). In the vocal folds specifically, implant formats provide mechanical stability and site retention, enabling therapy to remain in place long enough to support sustained treatment rather than relying on repeated injections (Cruz et al., 2025). Degradable polymer depots can shield therapeutics from rapid breakdown while enabling tunable, gradual release within the local microenvironment; by sustaining tissue drug levels, such systems may suppress fibrotic activity for longer than a transient injection (Awad et al., 2025). More generally, implants can release a drug in a controlled or sustained manner to maintain more constant drug exposure over time, which can reduce dosing frequency and minimize repeat procedures—an important practical advantage for a delicate tissue that is difficult to access repeatedly (Bhattacharjee et al., 2019; Huanbutta et al., 2024; L. Liu et al., 2017; Valvez et al., 2021). Finally, implants naturally enable hybrid delivery designs, where a biodegradable polymer implant serves as the long-residence “housing” while embedded nanoformulations provide controlled release—combining implant retention with nanoparticle-enabled tunability (Cruz et al., 2025; Manoukian et al., 2018; Padmasri & Nagaraju, 2021).

Although pirfenidone (PFD) has been explored as an antifibrotic candidate for vocal fold (VF) scarring in prior studies (Kodama et al., 2018b), most VF-focused efforts have evaluated (i) antifibrotic drugs, (ii) nanoparticle formulations, and (iii) implantable depots as separate strategies rather than as a single integrated platform. In parallel, irradiation-triggered VF therapy has been demonstrated using light-activatable local delivery systems, supporting the feasibility of spatially and temporally controlled dosing in the larynx (Zheng et al., 2025). However, few studies appear to combine PFD + biodegradable polymeric nanoparticles + an implant depot specifically for VF fibrosis, and reports of irradiation-triggered release from such PFD–nanoparticle–implant hybrid systems in VF models remain limited. Accordingly, this work addresses the gap by integrating PFD-loaded nanoparticles within a biodegradable implant to achieve sustained local exposure, while adding laser-triggered control to enable on-demand, dose-controllable release at the vocal folds. Together, sustained local exposure from the nanoparticle-in-implant platform and irradiation-enabled, on-demand control are positioned to reduce reliance on repeated vocal fold injections and extend therapeutic delivery at the site of fibrosis.

The objective of this study was to develop and evaluate a local, sustained, and dose-controllable pirfenidone (PFD) delivery platform for vocal fold applications by integrating PFD-loaded nanoparticles within biodegradable PLGA capsule implants. PFD was selected as a clinically established antifibrotic small molecule with a mechanism aligned to key profibrotic pathways implicated in vocal fold scarring. In this work, PFD-loaded nanoparticles were prepared and characterized using dynamic light scattering (DLS), transmission electron microscopy (TEM), and scanning electron microscopy (SEM), and then embedded into PLGA capsules designed for vocal fold placement. Baseline (unstimulated; 0-minute irradiation) release kinetics were quantified under physiological conditions, and the effect of laser irradiation (1- and 2-minute exposures) on PFD release was assessed relative to the unstimulated control to determine whether irradiation can provide on-demand enhancement of release from the hybrid implant system.

## 2. MATERIALS AND METHODS

### 2.1. Materials

Pirfenidone and poly(lactide-co-glycolide) (PLGA) (90:10; lactide:glycolide 90:10, MW 100,000–200,000; and 50:50; lactide:glycolide 50:50, MW 15,000–25,000) were purchased from PolyScitech. Gold nanorods (AuNRs) were obtained from NanoPartz, Inc. (Loveland, CO, USA). Phosphate-buffered saline (PBS; 10×) and 3.5 kDa MWCO dialysis membrane were purchased from Fisher Scientific. Dichloromethane (DCM) was purchased from Fisher Chemical. Poly (vinyl alcohol) (PVA; 98% hydrolyzed, average MW 13,000–23,000), 1.5 ml Eppendorf tube and UV-transparent microplates were purchased from Sigma-Aldrich.

### 2.2. PFD-Loaded PLGA nanoparticles

PFD–loaded PLGA nanoparticles were prepared using an oil-in-water (O/W) emulsion solvent-evaporation method. Briefly, PFD and PLGA were dissolved in dichloromethane (DCM, 4 mL) to form the organic phase. The aqueous phase consisted of poly (vinyl alcohol), dissolved in deionized (DI) water under stirring (350 rpm) for 1 h while heating at 90 °C until fully dissolved. The solution was then allowed to cool at room temperature. After cooling, the organic phase was added dropwise to the aqueous PVA solution in a 20 mL glass vial under stirring (350 rpm) over 10 min to form a pre-emulsion. The mixture was then probe-sonicated (20 kHz) using a pulsed program (10 s ON / 10 s OFF) with no ice to generate the emulsion. The resulting emulsion was stirred (350 rpm) at room temperature for 4 h to evaporate DCM. Nanoparticles were collected by centrifugation (5,000 × g, 10 min, 4 °C), washed twice with DI water to remove residual PVA and impurities, and resuspended in DI water (5 mL). The supernatant was used to determine the encapsulation efficiency. For long-term storage, nanoparticle suspensions were pre-frozen at −78 °C for 10 min and freeze-dried overnight using a freeze-dryer (Labconco, USA). The dried nanoparticles were collected in glass vials and stored at 4 °C until use.

#### 2.2.1. Optimization studies: size control

To obtain pirfenidone (PFD)–loaded PLGA nanoparticles with varied mean diameters, formulation and processing parameters were systematically varied, including PLGA mass (50–150 mg), PFD amount (20-50 mg), PVA concentration (0.5–1.5% w/v), and sonication time (1–2 min). Emulsification was performed using a probe sonicator operated at 20 kHz (40% amplitude; 500 W) without external cooling (no ice). Mean hydrodynamic diameter and polydispersity index (PDI) were measured by dynamic light scattering (DLS). For each optimization condition, entrapment efficiency (EE%) was determined indirectly by UV–Vis quantification of unencapsulated PFD in the supernatant (see section 2.3).

##### 2.2.1.1 Effects of PLGA amount on particle size

PLGA amount was varied (50–150 mg) while comparing matched PFD amount, PVA concentration, and emulsification conditions. The effect of PLGA content on nanoparticle properties was assessed using size, PDI, and EE%.

##### 2.2.1.2 Effects of PVA concentration on particle size

PVA concentration in the aqueous phase was varied (0.5–1.5% w/v) while comparing matched PLGA amount, PFD amount and emulsification conditions. Formulations were compared based on size, PDI, and EE%.

##### 2.2.1.3 Effects of sonication time on particle size

Sonication time was varied (1–2 min) under otherwise identical composition conditions and sonication power. Preliminary trials using longer sonication times were excluded from the final optimization because they produced visible aggregation and micrometer-sized particles.

Nanoparticles were evaluated by size, PDI, and EE% to identify conditions that reproducibly achieved the target size range.

##### 2.2.1.4 Effects of amount of PFD on particle size

PFD amount was varied between 20 and 50 mg while comparing matched PLGA amount, PVA concentration, and sonication time. The effects of PFD amount was evaluated based on particle size, PDI and EE% to determine whether higher drug loading altered nanoparticle size.

### 2.3. Characterization of PFD-Loaded PLGA nanoparticles

#### 2.3.1. Standard curve of PFD

Initially, standard curves of pirfenidone were obtained in phosphate buffer saline pH 7.4. Accurately weighed 10 mg of pirfenidone was transferred into a 10 ml volumetric flask, dissolved and diluted up to the mark to obtain a stock solution containing 1.0 mg/ml of pirfenidone. Aliquots from the stock solution were diluted with mobile phase to get the calibration standard solutions. Calibration standards were analyzed by UV–Visible spectrophotometer (SpectraMax M2e, USA) at 310 nm (λmax). SOFTMAX PRO 6.4.2 software was used for the Spectrophotometric analysis of pirfenidone samples. This study was conducted in triplicate, and the average of the three values was recorded. The calibration curve was made by plotting the absorbance of pirfenidone against respective concentrations. The calibration curve was constructed by plotting the peak area of pirfenidone against respective concentrations. It gives the absorption maxima (λmax) which were used to determine the amount of PFD in the in vitro dissolution sample and drug entrapped in a PLGA matrix.

#### 2.3.2. Dynamic Light Scattering (DLS): Measured mean particle size, polydispersity index (PDI), and zeta potential

The hydrodynamic size of the PLGA nanoparticles was determined using Dynamic Light Scattering (DLS) on a NanoBrook Omni particle analyzer (Brookhaven Instruments Co., Holtsville, NY, USA). The pirfenidone encapsulated PLGA nanoparticles were diluted 100-fold in a 10 mM KCl solution. Measurements were collected at a 90° scattering angle. For each sample, three consecutive 60-s acquisitions were recorded, and the entire measurement sequence was repeated twice to confirm reproducibility. Using the same instrument and dilution conditions, zeta potential was also measured based on electrophoretic mobility.

#### 2.3.3. Entrapment efficiency (EE%) and yield

Encapsulation efficiency was determined indirectly by quantifying the unencapsulated (free) pirfenidone present in the supernatant and wash fractions after nanoparticle purification. Briefly, after nanoparticle preparation, the suspension was centrifuged at 5,000 × g for 10 min at 4 °C and the supernatant was collected. The nanoparticle pellet was washed with DI water, and the wash solutions were also collected. The combined supernatant and wash fractions were analyzed using UV–Vis spectrophotometer (SpectraMax M2e, USA) at 310 nm (λmax) to quantify free PFD. The encapsulation efficiency was calculated from the difference between the total initial drug added and the free PFD measured in the supernatant/wash fractions according to the equation:

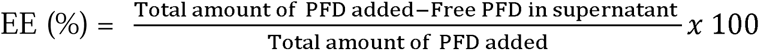

Nanoparticle yield was calculated by comparing the total recovered mass of formulated nanoparticles to the combined initial mass of polymer and pirfenidone used in the formulation. Measurements were performed in triplicate, and the mean yield was reported:

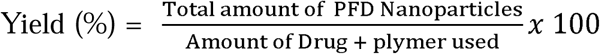

### 2.4. AuNR-coated PFD- loaded PLGA nanoparticles

PFD-loaded PLGA nanoparticles were prepared by an O/W emulsion solvent-evaporation method as described in Section 2.2.2. After EE determination, the nanoparticle suspension used for AuNR functionalization was further purified by dialysis using a 3.5 kDa molecular weight cutoff membrane overnight to remove any remaining free PFD, yielding a suspension containing drug-loaded nanoparticles. The dialyzed suspension was then centrifuged at 5,000 rpm for 1 h using an AccuSpin Micro 17 centrifuge (Fisher Scientific) to fractionate the formulation by size and isolate the larger nanoparticle fraction for AuNR functionalization. Citrate-coated gold nanorods (AuNRs; 10 nm diameter, 59 nm length; surface plasmon resonance peak ∼980 nm) were subsequently combined with the isolated nanoparticle fraction. AuNRs were added to the nanoparticle suspension at a 1:1 number ratio (AuNR:NP), calculated using the nanoparticle number concentration estimated from DLS-derived size measurements. The mixture was gently agitated overnight on an orbital shaker (New Brunswick Scientific) at 200 rpm to promote AuNR association with the nanoparticle surfaces via electrostatic interactions. The resulting AuNR–PFD–PLGA nanoparticle suspension was lyophilized overnight using a Labconco (USA) to obtain dry AuNR–PFD–PLGA nanoparticles for subsequent experiments.

### 2.5. Transmission Electron Microscopy (TEM) imaging

The morphology of the Naoparticles was characterized using a Talos L120C TEM (Thermo Fisher, Waltham, MA, USA). For TEM analysis, the PFD–PLGA (50:50) nanoparticles were examined. This nanoparticle suspension was then frozen at −80 °C for 1 h and subsequently lyophilized overnight with a freeze-dryer (Labconco, USA). This detailed TEM examination was critical for confirming the structural integrity of the nanoparticles, ensuring their suitability for further studies. For negative staining, a 2% (w/v) uranyl acetate (UA) solution was freshly prepared by dissolving 100 mg UA salt in 5 mL of boiling distilled water. The solution was stirred for 10 min in a covered beaker and then filtered into a labeled 5 mL black conical tube.

TEM grids were rendered hydrophilic using a glow discharger to improve wetting and staining. Two 50 μL droplets of the 2% UA solution were placed on parafilm, one serving as a wash droplet and the other as a stain droplet. The grid was first floated on the wash droplet for 10 s.

Next, a 3–4 μL aliquot of the sample (lyophilized nanoparticles resuspended and diluted) was applied to the grid and allowed to adsorb for 60 s. The grid was then transferred to the stain droplet and incubated for an additional 60 s. After each step, excess liquid was removed by gently tapping the grid against filter paper. Finally, the grid was dried using a microvacuum pump to remove residual liquid before imaging.

### 2.6. PFD release by NIR: Glass capillary experiment with/without AuNR

PFD-loaded PLGA nanoparticles and AuNR–PFD–loaded PLGA (50:50) nanoparticles were prepared to evaluate the effect of AuNR incorporation on PFD release under NIR irradiation. Approximately 30 μL of each nanoparticle suspension was introduced into a one-end-sealed glass capillary (average outer diameter: 1.27 mm; average wall thickness: 0.2 mm; Ei sco Laboratories, Victor) using a syringe fitted with an extended long needle (23 G × 3″, Air-Tite Products Co.). The loaded capillary was placed into a custom 3D-printed two-layer holder to maintain alignment with the incident laser beam. Irradiation was performed using a pulsed near-infrared (NIR) laser (WedgeHF, RPMC Lasers Inc.) operating at 1064 nm with a 10 kHz pulse repetition frequency and laser output at the fiber optic cable of 100 mW. The laser output was delivered through an optical fiber, which was inserted into the capillary to enable consistent irradiation along the full sample length. For each condition, the aqueous nanoparticle suspension (containing the same drug amount used in the corresponding implant release experiments) was exposed to irradiation durations of 1 or 2 min. Irradiation was applied in repeated 20 s ON / 40 s OFF cycles until the target total exposure time was reached. After irradiation, samples were recovered by opening the capillary ends, and a 20 μL aliquot was diluted 20-fold in PBS and incubated for 24 h. Following incubation, samples were processed using Amicon filters by centrifugation at 4000 × g for 10 min, and the filtrate was analyzed for PFD concentration by UV–Vis spectrophotometry at 310 nm using a pre-established calibration curve. Each condition was performed in triplicate, and total released PFD was quantified to determine the impact of AuNRs on nanoparticle drug release.

### 2.7. Implant fabrication and nanoparticle loading

The implants were produced following an established fabrication approach. In brief, PLGA was dissolved in dichloromethane (DCM) to prepare a 50 mg/mL polymer solution, and 1,500 μL of this solution was dispensed into a custom casting mold. The mold was made by supergluing two stainless-steel rectangular frames (4.9 cm × 3.1 cm × 1.5 cm; length × width × height) onto a phone glass protector. Solvent was then allowed to evaporate gradually overnight at 12 °C under a cover, yielding a solid polymer sheet. The dried sheet was removed with a razor blade, sectioned into 1 cm × 0.5 cm pieces, and each piece was rolled around a 22-gauge needle at 45 °C to form a double-layer tube (1 cm long; 0.0946 cm diameter). The resulting tube was cut into two 0.5 cm segments. One end of each segment was heat-sealed by clamping with a 60 °C iron. Each implant was then loaded with a pre-weighed mass of lyophilized PFD-loaded PLGA nanoparticle powder corresponding to ∼560 µg PFD per implant. Briefly, the implant was held vertically using tweezers to grip the sealed end at the top, with the open end oriented downward. The open end was brought into gentle contact with the nanoparticle powder and lightly tapped without applying force; with each tap, powder entered the lumen and progressively moved upward toward the sealed end. This was repeated until the entire pre-weighed powder mass was transferred into the implant. The open end was then heat-sealed, and both ends were reinforced with superglue.

The full implant fabrication workflow is summarized in Figure 1. The finished implants had final dimensions of 5 mm in length and 0.464 mm in outer diameter. The implant geometry was selected to fit within an 18-gauge syringe needle, enabling straightforward handling and placement. The total mass of each implant (capsule + contents) was approximately 2.06 mg, with the polymer capsule contributing ∼1.5 mg.

**Figure 1.**
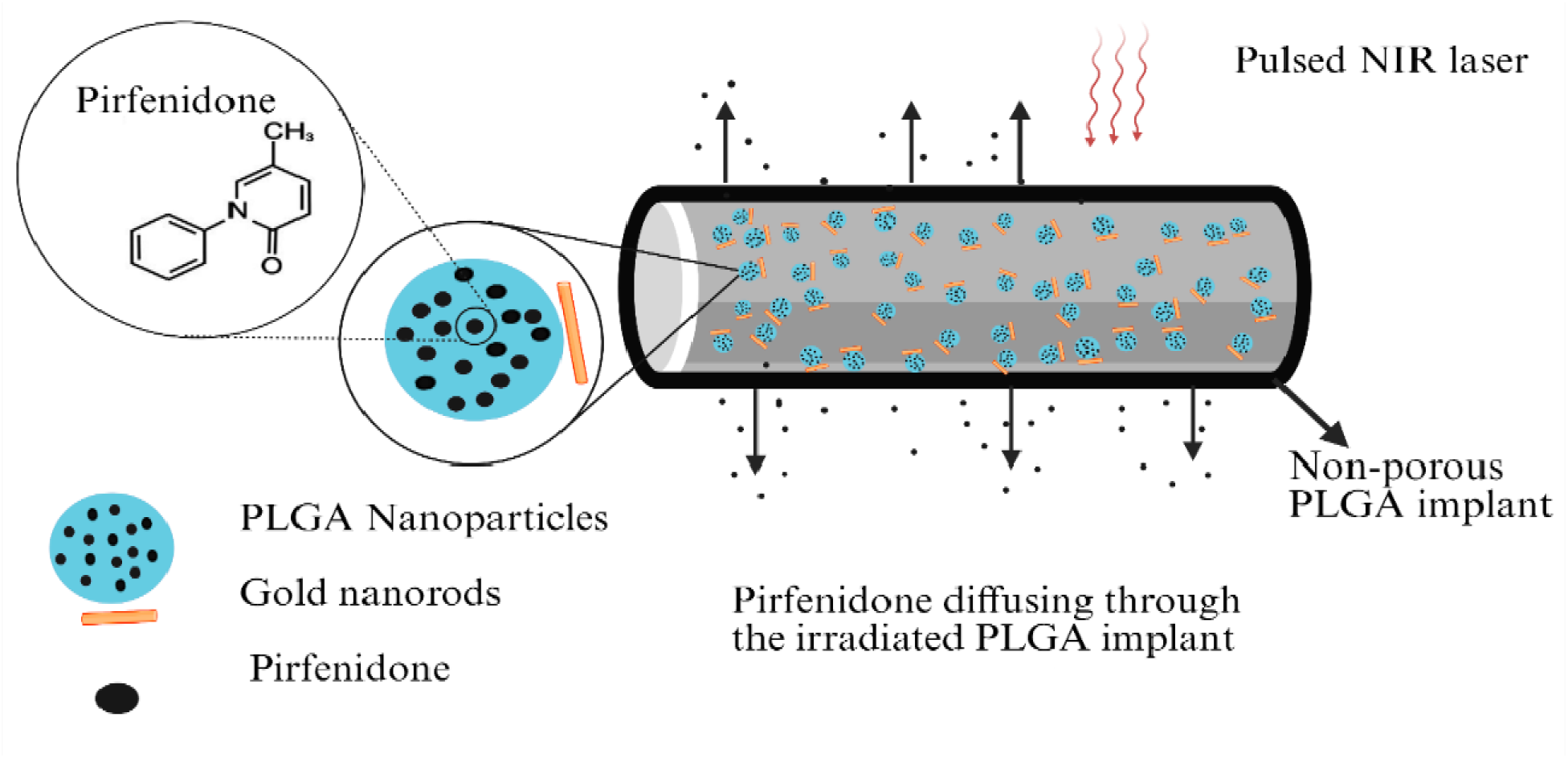
Schematic diagram of irradiation-enhanced pirfenidone release from non-porous PLGA implants. Under physiological conditions, a non-porous PLGA implant loaded with PFD-containing nanoparticles is designed to provide sustained baseline release. To test dose-controllability, brief NIR irradiation (1–2 min) is applied to determine whether it produces a measurable and repeatable increase in PFD release relative to the unstimulated (0-min) condition.

**Figure 2.**
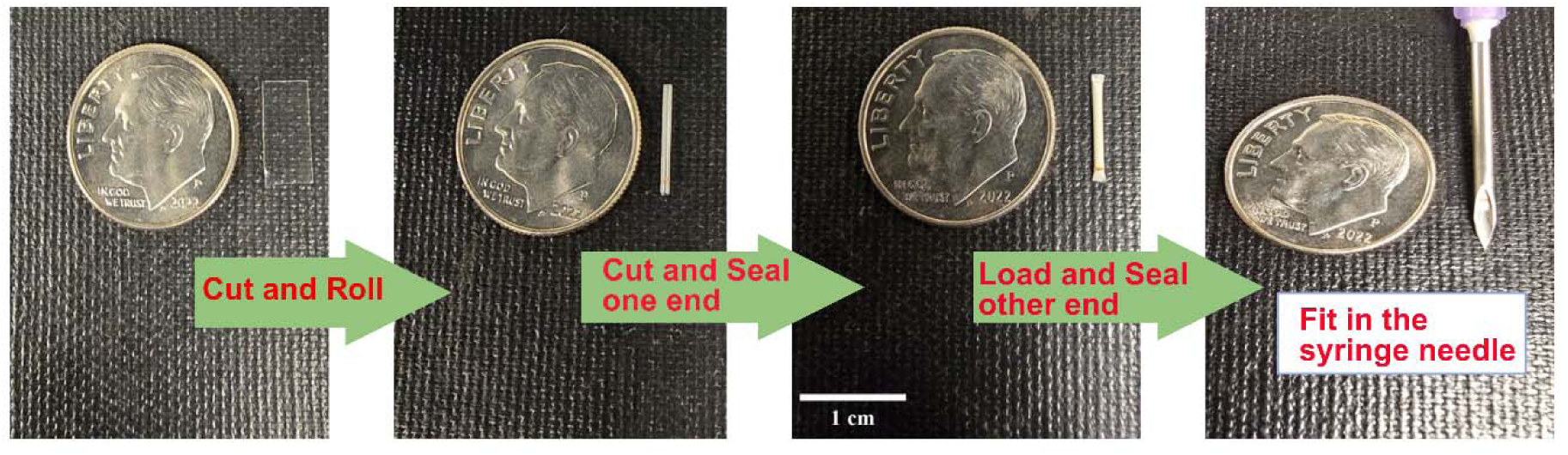
Fabrication process of PFD encapsulated nanoparticle PLGA implant (created based on (Zheng et al., 2023)

### 2.8. Scanning electron microscopy (SEM) imaging

SEM was used to examine both surface morphology and cross-sectional structure (Thermo Fisher Scios DualBeam SEM, Hillsboro, OR). Polymer sheet samples were mounted on either horizontal or vertical SEM stubs using double-sided graphite tape. Before imaging, samples were sputter-coated with an ∼5 nm Pt/Au layer for 10 s (Denton Vacuum Desk II, Moorestown, NJ).

To prepare cross-sections, 1 cm × 0.5 cm sheets were coated with glycerol, frozen in liquid nitrogen, and fractured to generate a fresh, clean break. The fractured specimens were rinsed with deionized water, dried overnight, mounted on vertical holders, and imaged to visualize the cross-section. Sheet thickness was quantified from SEM images using ImageJ (NIH).

### 2.9. In vitro NIR-triggered PFD release from the implant

Implants containing AuNR–PFD–loaded PLGA (50:50) nanoparticles were placed horizontally on a Petri dish to ensure uniform exposure during irradiation. A pulsed near-infrared (NIR) laser (WedgeHF, RPMC Lasers Inc.) operated at 1064 nm with a 10 kHz pulse repetition frequency and laser output at the fiber optic cable of 100 mW. Upon NIR exposure, AuNRs generate localized heat via plasmonic photothermal conversion, which increases polymer chain mobility and free volume in hydrated PLGA and thereby increases PFD diffusivity and permeability through the nanoparticle/implant matrix. Photothermal heating may also enhance water penetration and induce microstructural relaxation (e.g., transient pore formation), collectively facilitating irradiation-enhanced PFD transport compared with unstimulated conditions. For both the 1 min and 2 min irradiation durations, each implant received 5 s exposures at four distinct spots (totaling 20 s per cycle). Between cycles, a 40 s cooling period was implemented to limit overheating and reduce the risk of thermal damage to the polymer matrix. This cycle was repeated until the target total irradiation time was achieved. After irradiation, each implant was transferred into 1 mL PBS maintained at 35 °C in a 1.5 mL tube. PFD release was monitored by analyzing the PBS surrounding the implants. The PBS was replaced every 24 h during the first 2 weeks and then every weekday for the remainder of the study. Collected samples were quantified by UV–Vis spectroscopy at 310 nm, and cumulative PFD release was calculated from measured concentrations and plotted versus time to generate release profiles for each irradiation condition.

### 2.10. In vitro PLGA-PFD NP dialysis membrane release

3.5 kDa MWCO dialysis membrane was soaked in 1xPBS. 560 µg of pirfenidone entrapped in the PLGA nanoparticles were loaded into the dialysis membrane. The dialysis membrane loaded with nanoparticles was placed in a 1.5 ml Eppendorf tube with 1 ml of 1xPBS as a reservoir sink. The tubes were then incubated in a shaker at 35 °C. Every 24 hours the reservoir PBS was removed for UV-Vis analysis to determine and the pirfenidone concentration and the PBS was replaced with fresh PBS to maintain sink conditions. The release from the nanoparticles was determined over 40 days and compared to the release in the implants for the same duration.

## 3. RESULTS AND DISCUSSION

### 3.1 Formulation screening

Table 1 shows the effects of formulation and processing on the particle size, PDI and entrapment efficiency of PFD–PLGA nanoparticles prepared by the o/w emulsion-solvent evaporation method. Preliminary trials using a longer sonication time of 3-5 minutes produced visible aggregation and particle sizes in the micrometer range; therefore, these conditions were excluded from further optimization. The final screening focused on PLGA amount, PVA concentration, PFD amount, and sonication time using only 1 and 2 minutes.

**Table 1.**
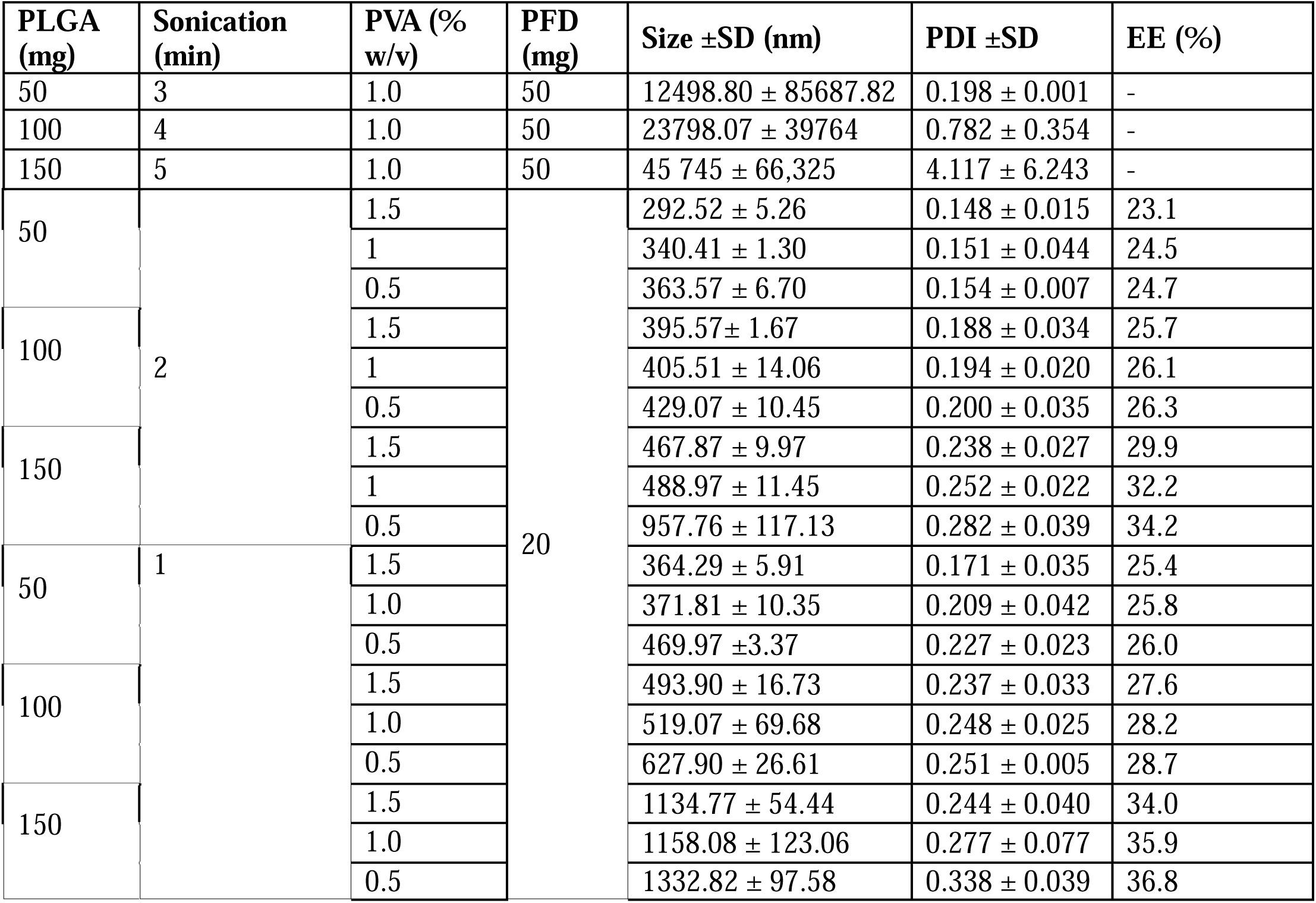
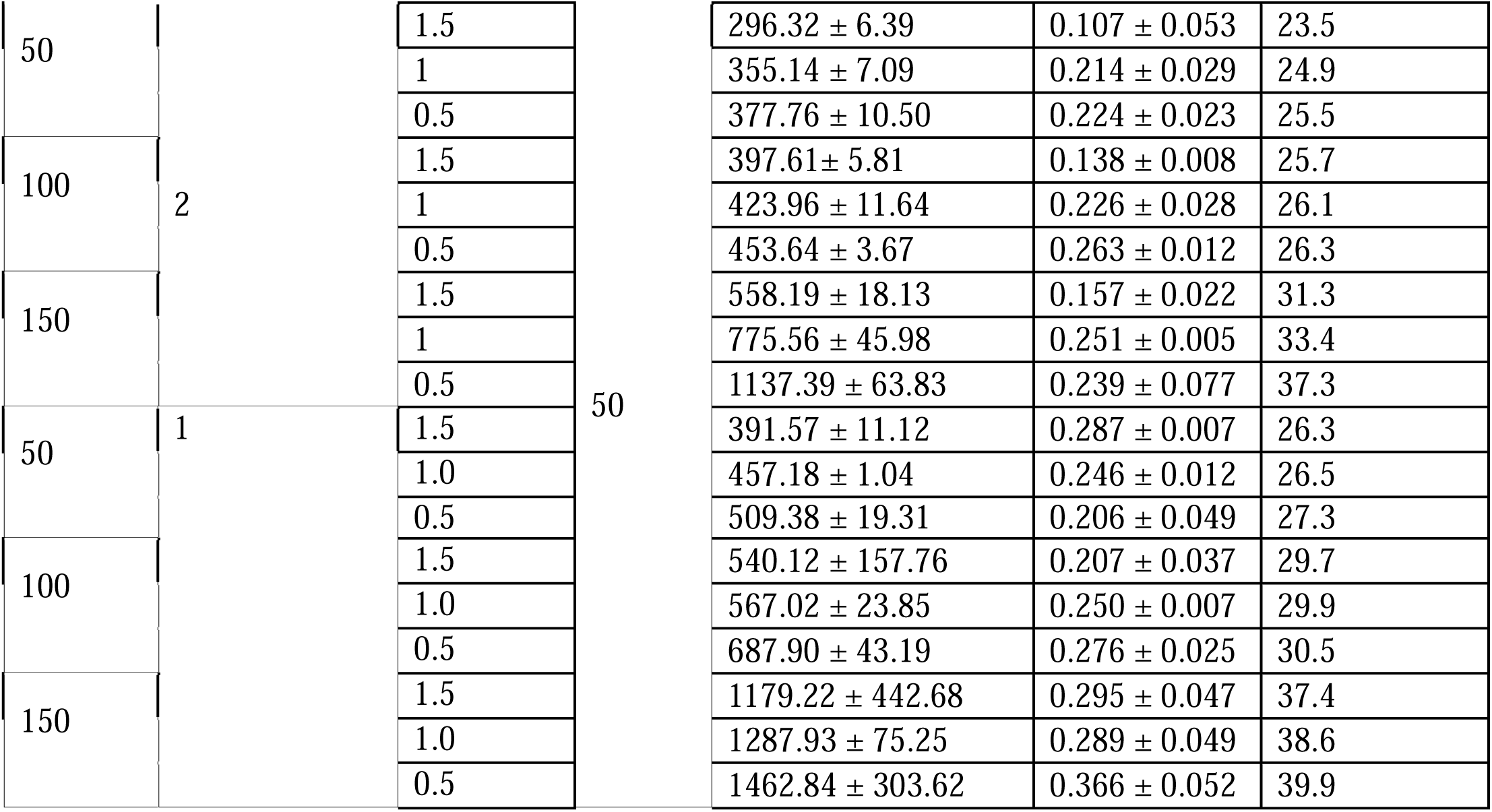
Effects of PLGA mass, sonication time, and PVA concentration on PFD–PLGA nanoparticle properties. Nanoparticles were prepared by an oil-in-water (o/w) emulsion–solvent evaporation method. Particle size (hydrodynamic diameter) and polydispersity index (PDI) were measured by DLS, and entrapment efficiency (EE) was determined as described in Methods.

#### 3.1.1. Effects of sonication time on particle size

Increasing probe sonication time from 1 to 2 min produced smaller nanoparticles with lower PDI values, indicating a narrower size distribution. Longer sonication provides greater emulsification energy, intensifying cavitation-driven shear, and promoting more complete breakup of the organic droplets during o/w emulsification (Muhaimin et al., 2025). As a result, smaller droplets are formed and are less likely to coalesce before hardening, yielding smaller and more uniform nanoparticles.

#### 3.1.2. Effect of PLGA amount on particle size

A progressive increase in particle size was observed as PLGA concentration increased from 50 to 150 mg. This trend is consistent with the viscosity–shear balance in emulsion-based nanoparticle formation: increasing polymer concentration raises the viscosity of the organic phase, increasing resistance to droplet deformation and reducing the effectiveness of shear-induced breakup during sonication (Tefas et al., 2015). Consequently, larger emulsion droplets persist and solidify into larger nanoparticles. A modest increase in PDI with PLGA mass also suggests slightly broader size distributions to higher polymer content. Entrapment efficiency increased with PLGA amount, likely due to a larger polymer matrix available for drug incorporation and reduced diffusion of the drug out of the organic droplets during hardening.

#### 3.1.3. Effect of PVA concentration on particle size

Increasing PVA concentration from 0.5% to 1.5%A generally decreases particle size and improved size distribution. PVA stabilizes the organic–aqueous interface and suppresses droplet coalescence, allowing smaller droplets generated during emulsification to remain stable long enough to harden into nanoparticles (Hernández-Giottonini et al., 2020). However, entrapment efficiency decreased with increasing PVA concentrations, which may be due to increased drug partitioning/solubilization into the continuous aqueous phase and/or greater loss during washing when stabilizer content is elevated (Kızılbey, 2019).

#### 3.1.4. Effect of PFD amount on particle size

Increasing PFD amount from 20 to 50 mg increased particle size across matched formulations Table 1). The increase is more pronounced at higher PLGA amounts and lower PVA concentrations, suggesting that higher drug loading contributed to larger particle size formation. The entrapment efficiency also increased with higher PFD loading, with the highest EE observed in larger particles, likely because larger particles provide greater polymer volume for drug retention (Salatin et al., 2017). Based on the screening results, three nanoparticle formulations were selected within the submicron range; the smallest (292nm), and the largest (957nm) were selected based on the particle size, while the medium-sized formulation (775 nm) was selected based on the higher entrapment efficiency, while remaining below 1000nm.

### 3.2 Characterization of the PFD-PLGA nanoparticle

DLS measurements were intensity-weighted, and the nanoparticle preparations were grouped into small, medium, and large batches based on their intensity-weighed mean hydrodynamic diameters. The small batch had an average hydrodynamic diameter of 292.52 ± 5.26 nm, the medium batch 775.56 ± 45.98 nm, and the large batch 957.76 ± 117.13 nm (Figure 3A–C).

**Figure 3:**
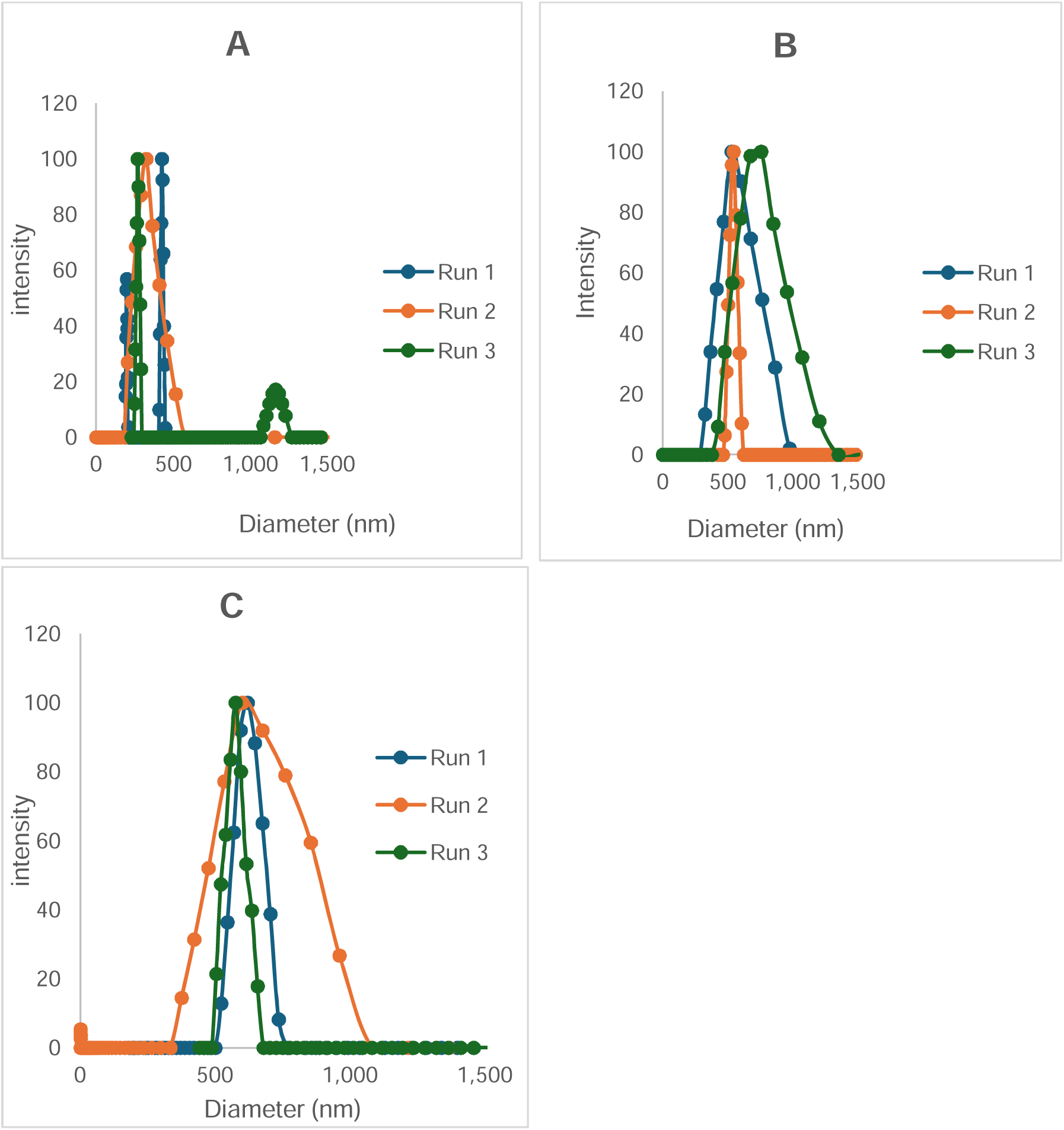
Intensity-weighted size distributions of PFD-loaded PLGA nanoparticles measured by DLS. (A) Small nanoparticle batch, (B) medium nanoparticle batch, and (C) large nanoparticle batch.

### 3.3. Characterization of the AuNR-coated PFD-PLGA nanoparticle

AuNRs were incorporated into the small-, medium-, and large-sized PFD-loaded PLGA nanoparticles, respectively. The attachment of AuNRs onto the surface of the PFD-loaded PLGA nanoparticles was confirmed by TEM imaging. Representative images of each group showed predominantly spherical nanoparticles and AuNR association with the nanoparticle surface (Figure 4A–C). The TEM micrographs are representative of the corresponding samples. The arrows indicate the AuNRs attached to the nanoparticle surface. Because the AuNR dimensions (10 nm × 59 nm) remained constant across all formulations, the AuNRs appeared relatively smaller as the nanoparticle size increased.

**Figure 4.**
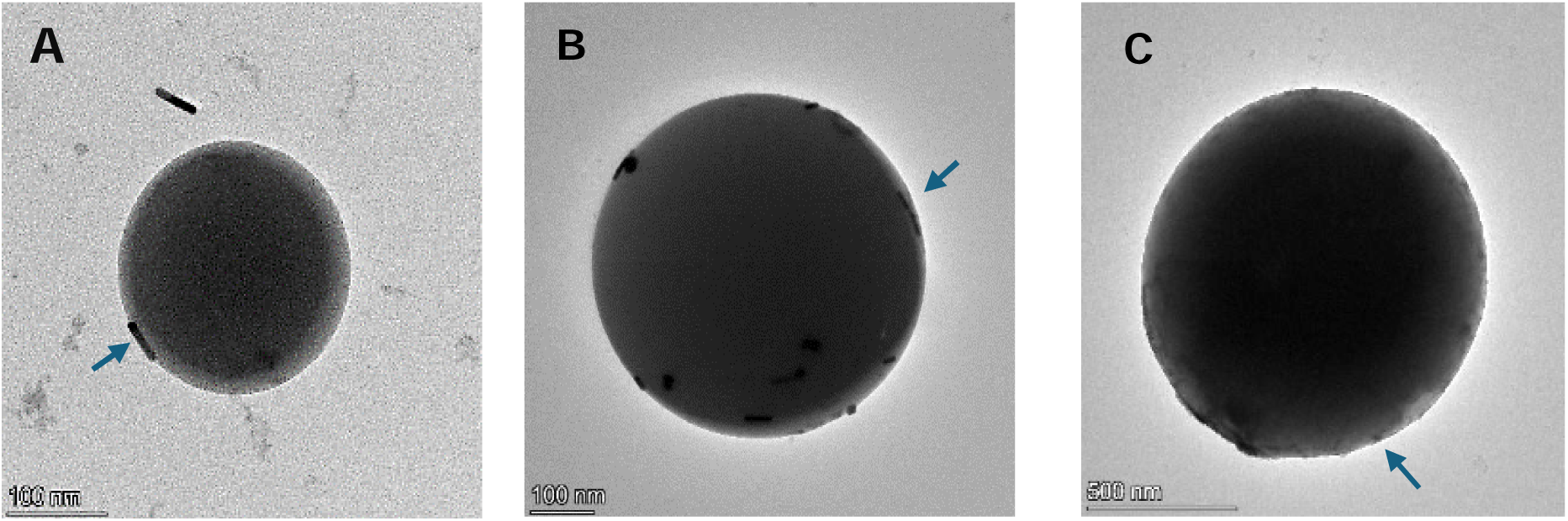
Representative TEM images of the small (A), medium (B), and large (C) nanoparticle batches.

### 3.3 Characterization of the PLGA capsule implant

The sheet top (Fig. 5A) appeared smooth, with no observable pores, characteristic of a homogenous surface with few visible bright particulates that could be debris. The bottom surface (Fig. 5B) was more heterogeneous, with small craters across the surface, indicating localized surface defects on the substrate-contact side. After rolling the sheet into an implant, the SEM image showing the outer surface together with the cross-section (Fig. 5C) indicates that the outer surface retains the crater texture observed on the substrate-facing side of the sheet, while the inner lumen surface appears comparatively smooth, except for a few visible artifacts, consistent with the air-facing layer forming the implant interior. A higher-magnification view of the outer surface (Fig. 5D) reveals a dense population of rounded craters with a partially patterned distribution across the implant exterior surface, with diameters ranging between 0.5-5.7 µm. The implant wall appears compact, with a dense and continuous cross-section across most of its thickness. The thickness of the PLGA capsule implant was analyzed using the cross-sectional (Figure 5E) image, and found to be 32.7 ± 3.9 µm. While craters are visible on the outer surface, there is no observable continuous, interconnected pore pathway extending from the outer surface through to the inner surface (Figure 5E). Rather, there are a few craters at the interface of the two layers that make the implant cross-section.

**Figure 5.**
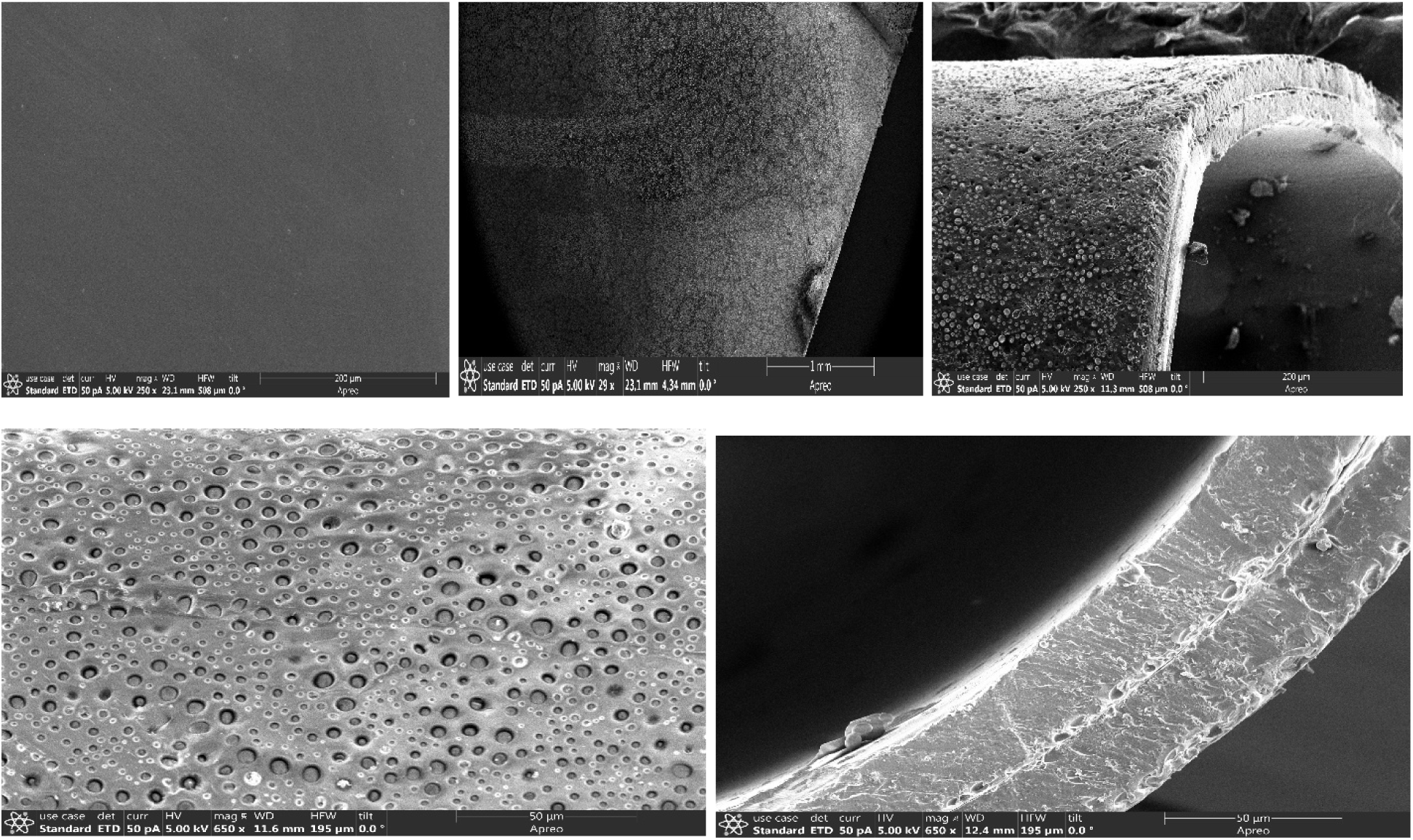
Scanning electron microscopy (SEM) images of PLGA (90:10) sheets and implants: (A) sheet, top surface (scale bar, 200 µm); (B) sheet, bottom surface (scale bar, 1 mm); (C) cross section (scale bar, 200 µm); (D) zoomed view of the implant top surface (scale bar, 50 µm); (E) implant cross-section (scale bar, 50 µm).

### 3.4. Early phase PFD release before implant breakage

In this study, the release was conducted for 278 days. However, the early phase refers to the first 110 days of release. This period was analyzed separately to better observe the effects of irradiation and nanoparticle size on PFD release, without the large increase in release caused by implant breakage at later time points.

#### 3.4.1. Effect of irradiation

The effects of irradiation on PFD release of different sizes PLGA nanoparticles *in vitro* are shown in Fig. 6. The release pattern of PFD from PLGA nanoparticles loaded in PLGA (90:10) implants was evaluated using PBS with a pH of 7.4 and temperature 37^0^ C. Overall, there was observed minimal burst release from all the samples, with a maximum burst 2.5% release of PFD achieved within 12 days.

**Figure 6.**
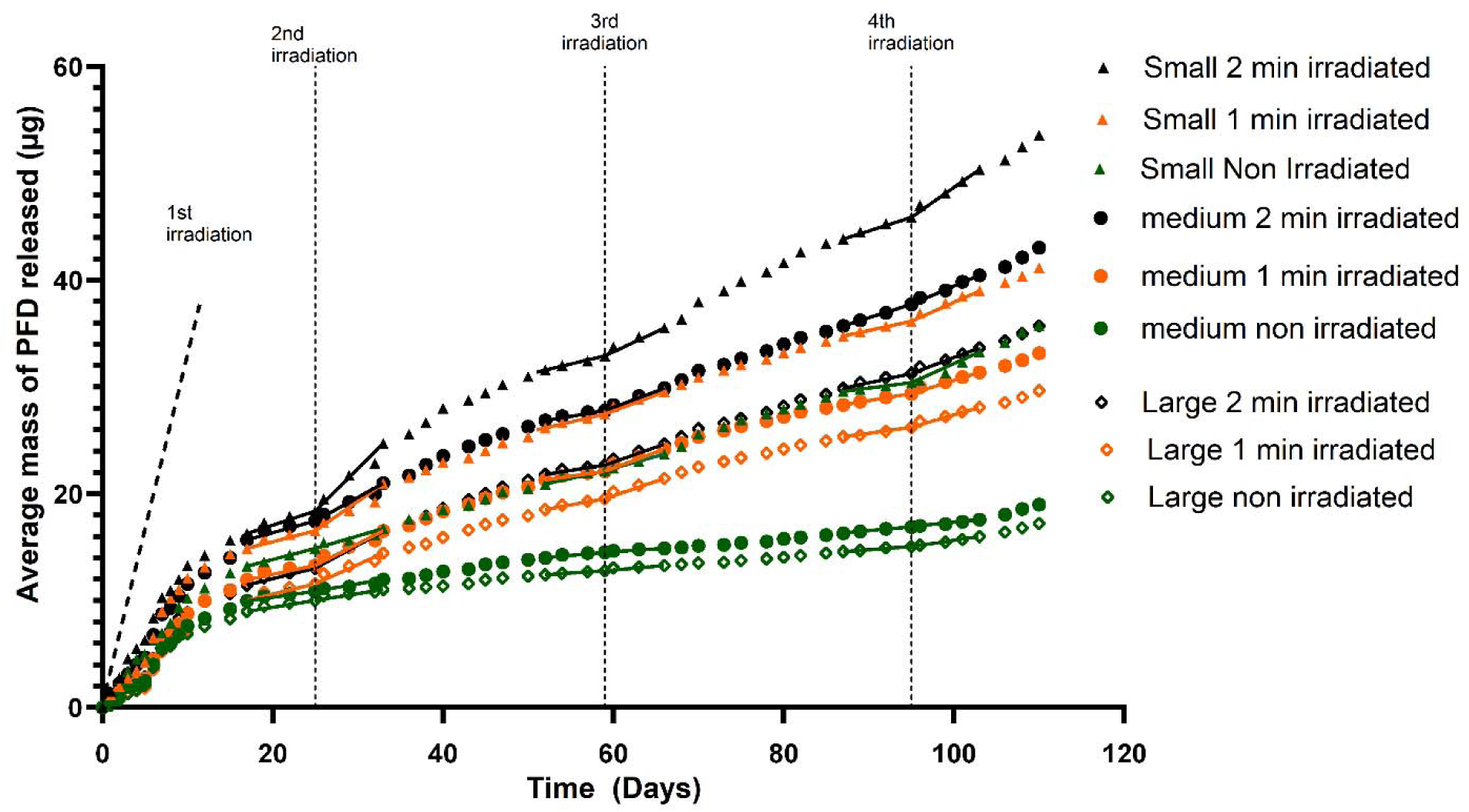
Average cumulative pirfenidone (PFD) release from PLGA 90:10 implants loaded with AuNR–PFD–PLGA nanoparticles. Each implant contained 560 µg PFD and was assigned to non-irradiated, 1-min irradiated, or 2-min irradiated groups. The graph shows the early release phase from Day 0 to Day 110 before implant breakage. Irradiation was applied on Days 0, 25, 59, and 95. Data are shown as mean ± SD, n = 3.

All three groups of nanoparticles exhibited increased PFD release after irradiation compared to controls. The higher levels of PFD release after irradiation could be attributed to the photothermal effects of the AuNRs. Gold Nanorods (AuNR) are capable of absorbing near-infrared light and converting it to heat through their plasmonic properties (Zhou et al., 2022). As a result of the localized heating produced by the AuNRs, chain mobility of the polymers increased, permitting PFD to diffuse more readily from within the nanoparticle-loaded implant.

Four irradiation events were applied during the release study, and the dashed vertical lines in Figure 6 indicate the irradiation time points. Changes in the release slope (lines) following each irradiation suggest irradiation-enhanced PFD release. To assess the effects of irradiation more directly, the change in the gradient of release pre-irradiation and post-irradiation was determined (Table S1, Supplementary). For the small nanoparticle group, the irradiation effect was strongest in the 2-min irradiated group, especially after the 2nd irradiation. The release gradient increased by +0.533 µg/day in the 2-min group, compared with +0.347 µg/day in the 1-min group and only +0.025 µg/day in the non-irradiated group. This suggests that longer irradiation produced a stronger increase in PFD release rate for the small nanoparticle implants. A similar trend was observed in large-sized nanoparticles. For the medium nanoparticle group, both irradiated groups showed increased release rates compared with the non-irradiated control. However, the 1-min irradiated group showed a slightly higher change in gradient than the 2-min group. This suggests that irradiation enhanced release in the medium group, but the response did not increase strictly with irradiation duration.

#### 3.4.2 Effect of size of PLGA-nanoparticles on PFD release

Figure 6 also shows the effect of PLGA nanoparticle size on early-phase PFD release. To evaluate the role of nanoparticle size, the non-irradiated groups were first compared because they represent passive release without additional effects of irradiation. By day 110, the small, medium and large nanoparticles -loaded implants cumulatively released 38.3 µg, 21.1 µg and 18.9 µg of PFD, respectively, confirming the trend small> medium.> large. A similar size-dependent trend was observed in the irradiated groups, where the small, medium and large nanoparticles -loaded implants cumulatively released 44.4 µg, 36.5 µg, and 32.1 µg in the 1 min irradiated groups and 56.7 µg, 46.2 µg, and 38.3 µg in the 2 min irradiated groups. The higher release from smaller nanoparticles may be attributed to their larger surface-area-to-volume ratio and shorter diffusion path length, which allow PFD to diffuse out more readily before passing through the surrounding PLGA implant matrix (Siepmann et al., 2004).

### 3.5 Long-term PFD kinetics

#### 3.5.1 Small-sized PLGA nanoparticle group

PFD release from Day 0 to Day 197 increased gradually, as shown in Fig 7. In this study period, the 2 min irradiated group had the highest PFD release, followed by the 1-min group, while the non-irradiated group showed the lowest release. This release order indicates that laser irradiation enhanced PFD release. The 2 min group released 113 µg before implant fractured on day 197. In the late phase (beyond day 197-199), there is a sharp increase in the PFD released, with the release no longer following small > medium > large. This is because of implant breakage, releasing nanoparticles and PFD that were trapped within the implant.

**Figure 7:**
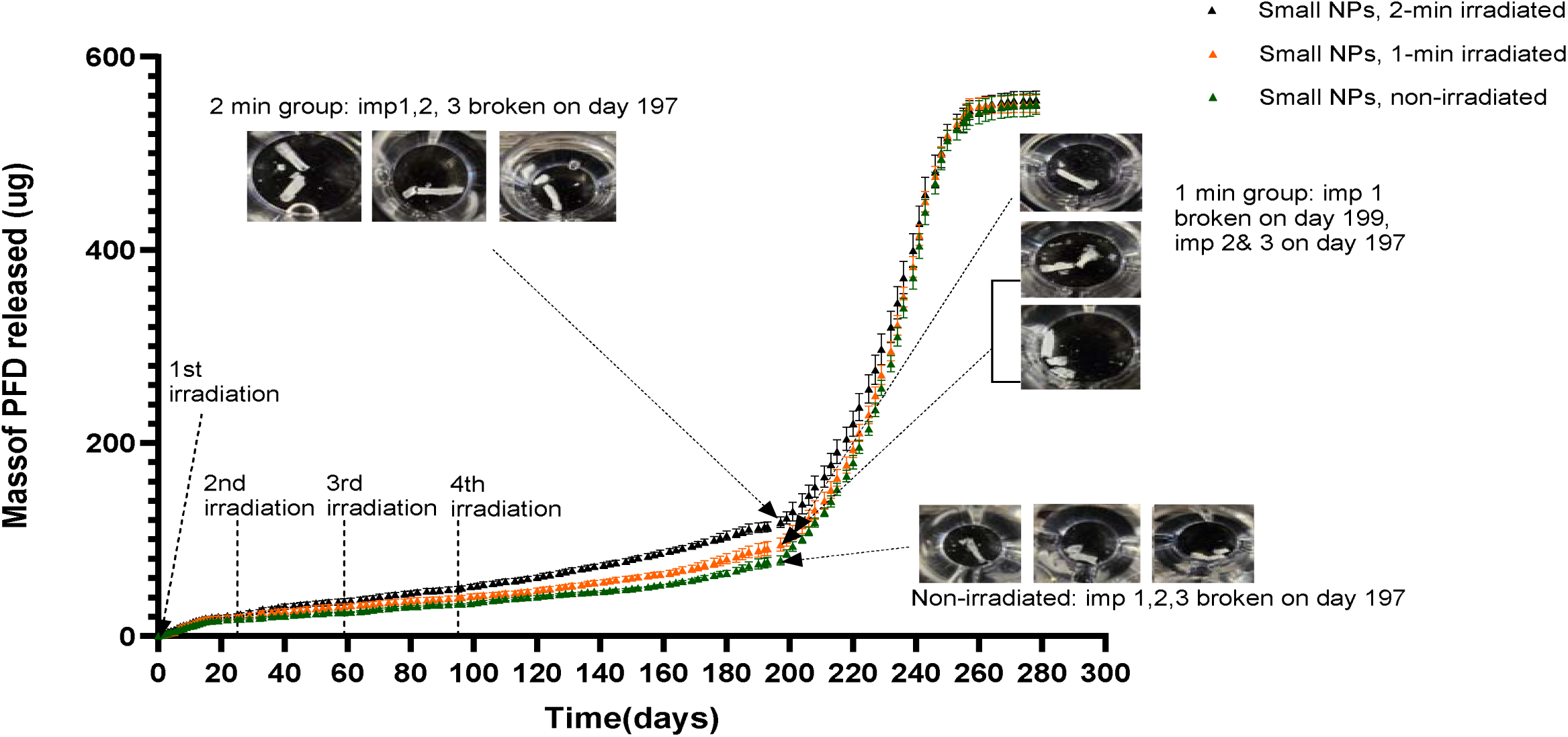
Average cumulative pirfenidone (PFD) release from PLGA (90:10) implants (0.9 mm diameter × 5 mm length) loaded with small-sized AuNR–PFD nanoparticles, corresponding to 560 µg PFD per implant, irradiated for 0 min, 1 min, or 2 min. Error bars represent standard deviation across three implants (n = 3), each measured in triplicate via UV–Vis spectrophotometry. Irradiation was applied on Day 25, 59 and 95 for the 1-min and 2-min groups.

**Figure 8:**
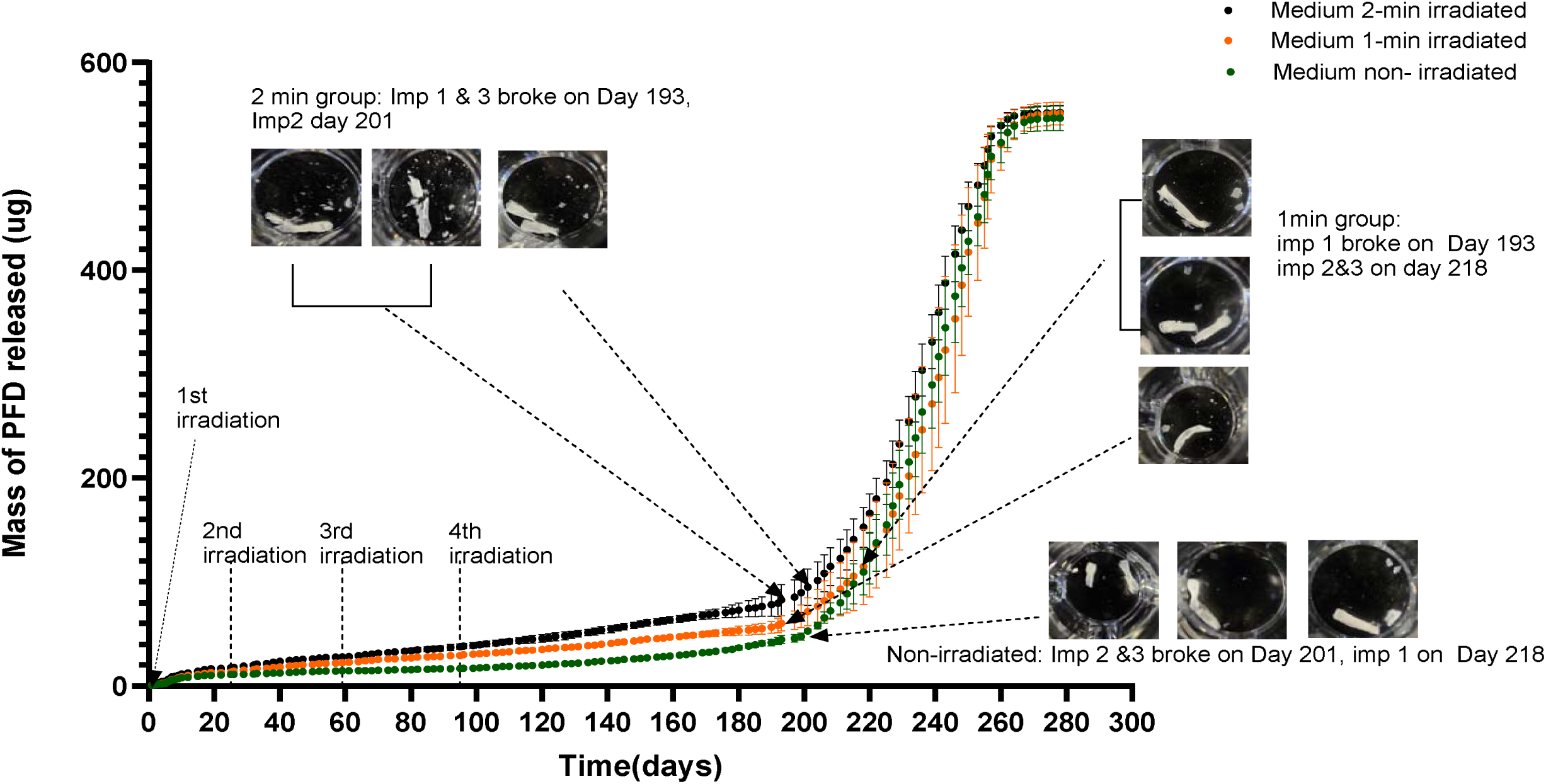
Average cumulative pirfenidone (PFD) release from PLGA (90:10) implants (0.9 mm diameter × 5 mm length) loaded with medium-sized AuNR–PFD nanoparticles, corresponding to 560 µg PFD per implant, irradiated for 0 min, 1 min, or 2 min. Error bars represent standard deviation across three implants (n = 3), each measured in triplicate via UV–Vis spectrophotometry. Irradiation was applied on Day 25, 59 and 95 for the 1-min and 2-min groups.

The slow drug-release mechanism can be understood in terms of bulk erosion of PLGA. Upon water imbibition, PFD located near the surface is easily dissolved and diffuses out (Rodrigues De Azevedo et al., 2017), explaining why all groups showed an initial increase in cumulative PFD release. Absorbed water causes hydrolysis degradation of the polymer chains into mono- and oligomeric acids (Klose et al., 2006). These generated acids diffuse into the bulk fluid and are neutralized, while bases from the bulk fluid diffuse into the nanoparticles. However, the diffusion rate of bases into the nanoparticles is slower than the rate of acid generation. Thus, pH in the nanoparticle core decreases due to acid accumulation, accelerating polymer degradation since protons catalyze ester bond cleavage (Klose et al., 2006). This releases PFD from the degrading nanoparticles into the implant lumen, from which it is released slowly. Since PLGA (90:10) implant degradation is much slower than PLGA (50:50) nanoparticle degradation, the implant acts as a long-term barrier and delays the rapid loss of nanoparticle-associated PFD.

Pirfenidone is relatively hydrophobic, and the PLGA (90:10) matrix is predominantly hydrophobic, which favors drug retention within the polymer phase, accounting for the slow release observed in the early and intermediate phase. However, once the implant fractured, previously internal regions containing partially degraded PLGA (50:50) nanoparticles became directly exposed to the aqueous medium. This structural disruption increased the effective surface area, shortened diffusion pathways, and allowed both nanoparticle and PFD to escape.

#### 3.5.2 Medium-sized PLGA nanoparticle batch

Similar to the small-sized group, PFD release before breakage also showed a slow and sustained increase in the early and intermediate phase, with the irradiated groups releasing more PFD than the non-irradiated group. Compared with the small-sized batch, the 2 min group released 85 µg before implant fracture on day 193 (Fig 7). This could be because the medium-sized nanoparticles may have longer diffusion pathways, which can reduce the rate at which PFD is released from the nanoparticles before implant fracture.

#### 3.5.3 Large size PLGA nanoparticle batch

A release trend similar to that observed for the small- and medium-sized nanoparticle groups was observed for the large-sized nanoparticle group over the early and intermediate release phase. PFD release remained gradual and sustained, with the 2-min and 1-min irradiated groups releasing more PFD than the non-irradiated group, indicating that laser irradiation enhanced drug release while the implant remained intact. However, the large-sized nanoparticle group irradiated for 2 min released only 71.4 µg of PFD by approximately Day 201 (Fig 9), which was lower than the 2-min irradiated small- and medium-sized nanoparticle groups. This lower release may be due to the longer internal diffusion pathways and lower surface-area-to-volume ratio of the larger nanoparticles.

**Figure 9:**
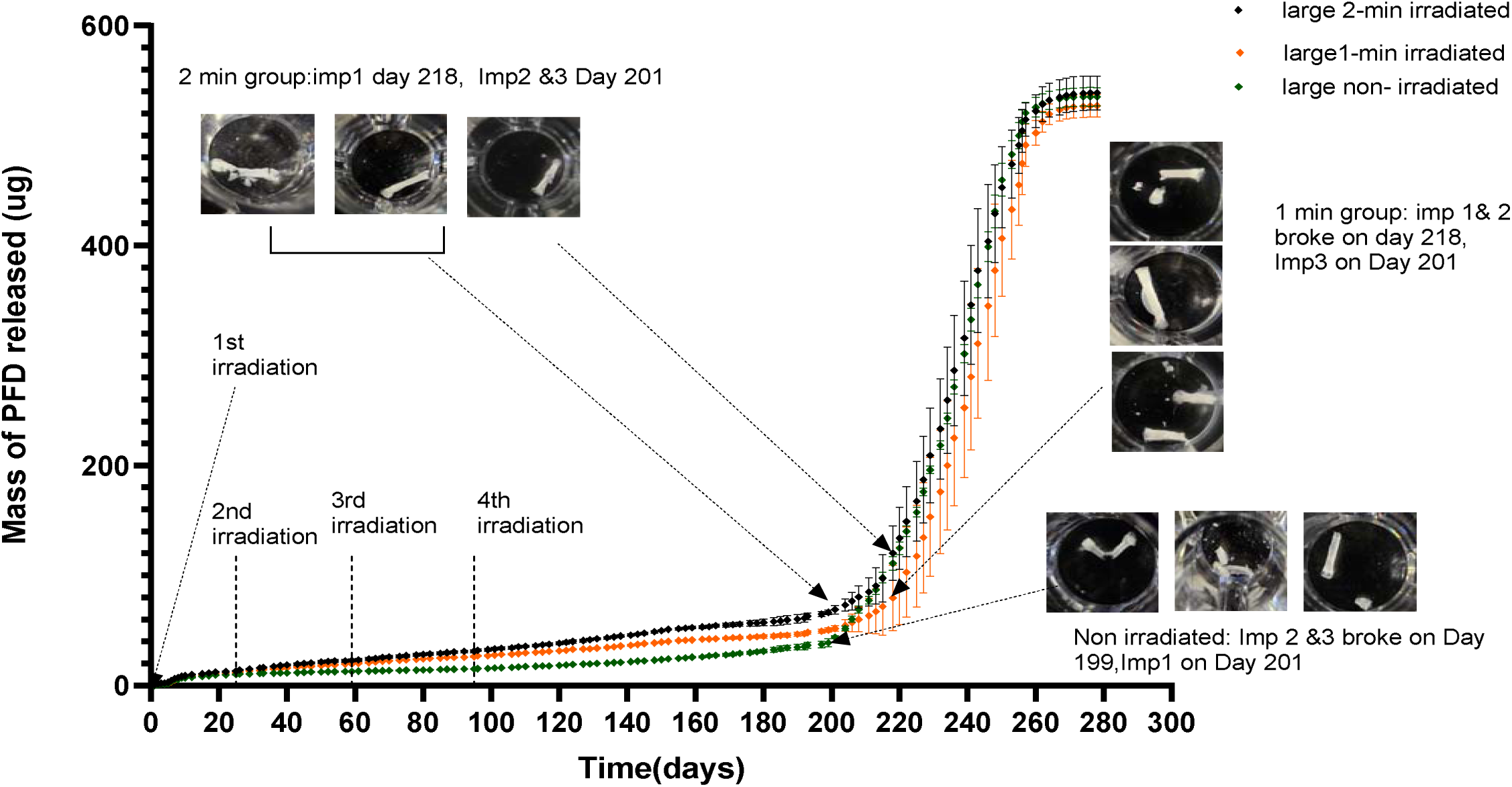
Average cumulative pirfenidone (PFD) release from PLGA (90:10) implants (0.9 mm diameter × 5 mm length) loaded with large-sized AuNR–PFD nanoparticles, corresponding to 560 µg PFD per implant, irradiated for 0 min, 1 min, or 2 min. Error bars represent standard deviation across three implants (n = 3), each measured in triplicate via UV–Vis spectrophotometry. Irradiation was applied on Day 25, 59 and 95 for the 1-min and 2-min groups.

PFD release from nanoparticle-loaded PLGA implants reached only approximately 20% of the initial 560 µg PFD loading after approximately 190 days. Table 2 summarizes the amount of PFD released by each implant immediately before implant fracture.

**Table 2.** Percentage of initial PFD load released before implant fracture from small, medium, and large nanoparticle-loaded PLGA implants.

| *Nanoparticle size* | Small size-Non irradiated | Small size 1-min irradiated | Small size 2-min irradiated | Medium size non-irradiated | Medium size 1 min irradiated | Medium size 2 min irradiated | Large size non irradiated | Large size 1 min irradiated | Large size 2 min irradiated |
| --- | --- | --- | --- | --- | --- | --- | --- | --- | --- |
| *% load released* | 13.7 | 16.2 | 20.2 | 8.8 | 11.0 | 14.8 | 6.9 | 9.4 | 12.3 |

The findings for the drug release from small, medium, and large particle size nanoparticles indicate a mismatch between intended drug release and the implant degradation, with the implants degrading after approximately 190 days. The sharp increase after implant breakage could lead to drug dumping, causing toxicity. Indicating the need for future formulation that would maintain integrity throughout the treatment period, preventing the rapid release observed after implant fracture.

### 3.6 Dialysis Membrane Release

To determine the cause of the slow release, small, medium and large nanoparticles formulated from PLGA 50:50 encapsulating 560 µg PFD were further evaluated using the dialysis bag method. For the experimental setup, the dialysis bag loaded with nanoparticles was placed in a 1.5 ml tube containing 1ml PBS at pH 7.4. The tube was placed in the agitator. PBS permeates through the dialysis membrane and the nanoparticles, dissolves the drug, and permeates from the dialysis bag into the bulk solution (Yu et al., 2019). The PBS in the receiver compartment was analyzed using UV after 24 hours and replaced with fresh PBS to maintain sink conditions. The in vitro drug release kinetics obtained were compared with those of PFD released from nanoparticles loaded in an implant, and the results are presented in Figure 10.

**Figure 10.**
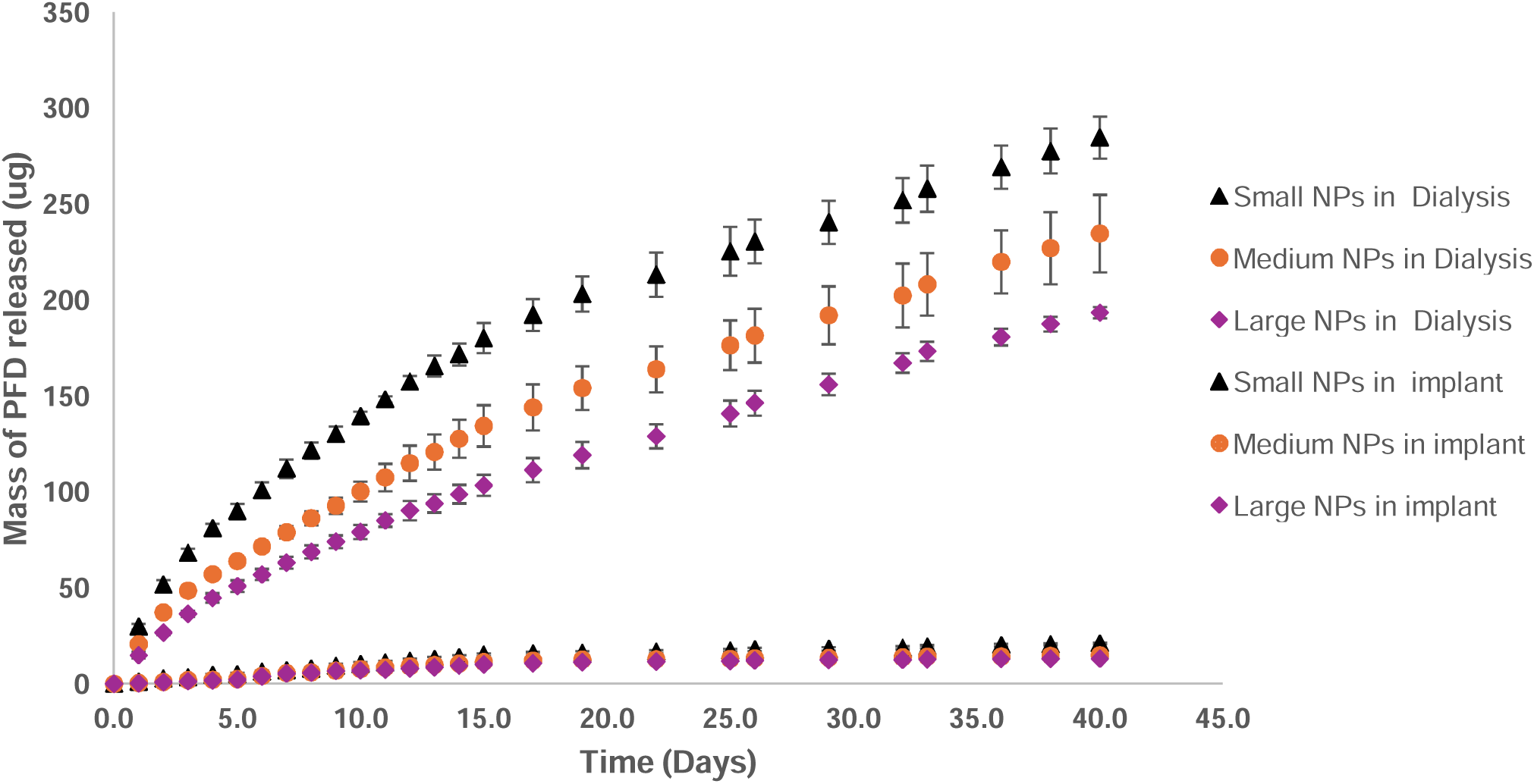
Comparison of average cumulative PFD release from PFD-loaded PLGA nanoparticles loaded in PLGA (90:10) implants and 3.5 kDa MWCO dialysis membranes for small, medium and large nanoparticles. PLGA (90:10) implants (0.9 mm diameter × 5 mm length) and 3.5 kDa MWCO dialysis membranes were each loaded with nanoparticles corresponding to 560 µg PFD.

Approximately 50.8% of the initial loading of PFD was released from small-sized nanoparticles compared to 3.8% released from implants over the same study duration. Similarly, for the medium-sized and large-sized nanoparticles, 41.9% and 34.5% were released from the dialysis bag, and approximately 2.7% and 2.3%, respectively, from implants. This indicates that when the nanoparticles are not surrounded by the PLGA matrix, PFD diffuses more readily, indicating that the PLGA implant acts as an additional diffusion barrier. A size-dependent release trend from the dialysis group, with small nanoparticles exhibiting the highest PFD release, followed by medium and large nanoparticles. This is consistent with a large surface area to volume ratio. Conversely, a similar trend was observed in the release of PFD from the nanoparticles loaded in the implant.

### 3.7 PFD release kinetics: PLGA nanoparticles strategy to modulate the implants

The mismatch between sustained PFD release and implant degradation, resulting in an abrupt increase in PFD release, suggests the need for better alignment of drug diffusion with polymer breakdown. The release of PFD beyond 6 months, a slower-degrading polymer than PLGA 90:10, could be considered. Compared to PLGA 90:10, PCL degrades more slowly due to its semicrystalline and hydrophobic nature, with a reported degradation time of approximately 2-4 years (Dias et al., 2022). In contrast, PLGA degradation is influenced by the lactide: glycolide ratio. The higher the glycolide ratio, the higher the water uptake, increasing the degradation rate. In the present study, though PLGA 90:10 degraded slowly, compared to PCL, the degradation was substantially faster. For a shorter duration release of PFD, the strategy should focus on increasing the permeability of the PLGA 90:10 with the incorporation of a porogen such as PEG. As PEG dissolves in an aqueous release medium, it creates a porous PLGA matrix, allowing more water penetration, increased contact surface, and improved drug diffusion. Implant wall thickness can be adjusted to reduce the diffusion barrier. In this system, the wall thickness could be adjusted by reducing the polymer casting volume or increasing the casting area. A thinner wall could shorten the diffusion path between the nanoparticle-loaded lumen and the external medium, allowing more release of PFD before the structural breakdown.

## Conclusion

In this study, a biodegradable nanoparticle-in-implant delivery platform was successfully developed for sustained and light-facilitated pirfenidone (PFD) release for potential vocal fold fibrosis applications. PFD-loaded PLGA nanoparticles with varying particle sizes were incorporated into PLGA implants, and AuNR integration enabled irradiation-enhanced drug release under near-infrared (NIR) exposure. The implants demonstrated prolonged PFD release for more than 190 days with minimal burst release, while laser irradiation increased release rates compared with non-irradiated controls. Nanoparticle size significantly influenced release behavior, with smaller nanoparticles exhibiting greater cumulative release due to shorter diffusion pathways and higher surface-area-to-volume ratios. Dialysis membrane studies further confirmed that the PLGA implant acted as an additional diffusion barrier that substantially prolonged release kinetics. Although late-stage implant fracture resulted in accelerated drug release, the findings demonstrate the feasibility of combining sustained local delivery with externally controllable release enhancement in a single hybrid platform. Overall, this system shows promise as a long-acting and dose-controllable local antifibrotic therapy for vocal fold fibrosis and may be adaptable to other localized fibrotic disorders requiring prolonged therapeutic delivery.

