## Supplementary material for "Biodegradable Nanoparticle-in-Implant Platform for Sustained and Light-Boosted Pirfenidone Delivery": Table S1

**Supplementary information**

**Table S1. Pre- and post-irradiation PFD release gradients during the early release phase**

| **Size** | **Condition** | **Pre-2nd (µg/day)** | **Post-2nd (µg/day)** | **Δ2nd (µg/day)** | **Pre-3rd (µg/day)** | **Post-3rd (µg/day)** | **Δ3rd (µg/day)** | **Pre-4th (µg/day)** | **Post-4th (µg/day)** | **Δ4th (µg/day)** |
| --- | --- | --- | --- | --- | --- | --- | --- | --- | --- | --- |
| **Small** | **0 min** | 0.208 | 0.233 | +0.025 | 0.175 | 0.225 | +0.050 | 0.101 | 0.359 | +0.258 |
| **Small** | **1 min** | 0.205 | 0.552 | +0.347 | 0.185 | 0.300 | +0.115 | 0.174 | 0.354 | +0.180 |
| **Small** | **2 min** | 0.260 | 0.793 | +0.533 | 0.191 | 0.370 | +0.179 | 0.254 | 0.560 | +0.306 |
| **Medium** | **0 min** | 0.112 | 0.140 | +0.028 | 0.073 | 0.053 | −0.020 | 0.074 | 0.081 | +0.007 |
| **Medium** | **1 min** | 0.165 | 0.405 | +0.240 | 0.117 | 0.295 | +0.178 | 0.133 | 0.248 | +0.115 |
| **Medium** | **2 min** | 0.219 | 0.442 | +0.223 | 0.133 | 0.299 | +0.166 | 0.252 | 0.341 | +0.089 |
| **Large** | **0 min** | 0.129 | 0.128 | −0.001 | 0.052 | 0.069 | +0.017 | 0.060 | 0.118 | +0.058 |
| **Large** | **1 min** | 0.193 | 0.358 | +0.165 | 0.144 | 0.271 | +0.127 | 0.110 | 0.233 | +0.123 |
| **Large** | **2 min** | 0.198 | 0.431 | +0.233 | 0.124 | 0.280 | +0.156 | 0.174 | 0.300 | +0.126 |
